## Supplementary Materials for "Adversarial attacks and adversarial robustness in computational pathology"

### Supplementary Tables

| Study design (Part 1) | Completed, page | |
|---|---|---|
| The clinical problem in which the model will be employed is clearly detailed in the paper. | yes | # |
| The research question is clearly stated. | yes | # |
| The characteristics of the cohorts (training and test sets) are detailed in the text. | yes | # |
| The cohorts (training and test sets) are shown to be representative of real-world clinical settings. | yes | # |
| The state-of-the-art solution used as a baseline for comparison has been identified and detailed. | yes | # |
| **Data and optimization (Parts 2, 3)** | | |
| The origin of the data is described and the original format is detailed in the paper. | yes | # |
| Transformations of the data before it is applied to the proposed model are described. | yes | # |
| The independence between training and test sets has been proven in the paper. | yes | # |
| Details on the models that were evaluated and the code developed to select the best model are provided. | yes | # |
| Is the input data type structured or unstructured?: **Unstructured images** | | |
| **Model performance (Part 4)** | | |
| The primary metric selected to evaluate algorithm performance (e.g., AUC, F-score, etc.), including the justification for selection, has been clearly stated. | yes | # |
| The primary metric selected to evaluate the clinical utility of the model (e.g., PPV, NNT, etc.), including the justification for selection, has been clearly stated. | yes | # |
| The performance comparison between baseline and proposed model is presented with the appropriate statistical significance. | yes | # |
| **Model examination (Part 5)** | | |
| Examination technique 1a: **Highly scoring tiles (qualitative and quantitative analysis)** | yes | # |
| Examination technique 2a: **Principal component analysis (PCA)** | yes | # |
| A discussion of the relevance of the examination results with respect to model/algorithm performance is presented. | yes | # |
| A discussion of the feasibility and significance of model interpretability at the case level if examination methods are uninterpretable is presented. | yes | # |
| A discussion of the reliability and robustness of the model as the underlying data distribution shifts is included. | yes | # |
| **Reproducibility (Part 6)** | | |
| Tier 1: complete sharing of the code | yes | # |

**Suppl. Table 1: Minimum information about clinical artificial intelligence modeling (MI-CLAIM) checklist**

| Task | Run | Normal model | | Adversarially trained model | | |
|---|---|---|---|---|---|---|
| | | ResNet | ViT | ResNet | ResNet with DBN | ViT |
| **RCC subtyping** *raw data for Figure 1C* | AUROC of run #1 | 0.967 [0.953 - 0.978] | 0.965 [0.952 - 0.976] | 0.956 [0.942 - 0.969] | 0.902 [0.880 - 0.923] | 0.940 [0.923 - 0.957] |
| | AUROC of run #2 | 0.975 [0.964 - 0.984] | 0.962 [0.949 - 0.973] | 0.958 [0.944 - 0.971] | 0.947 [0.931 - 0.962] | 0.941 [0.924 - 0.958] |
| | AUROC of run #3 | 0.948 [0.931 - 0.965] | 0.941 [0.923- 0.958] | 0.956 [0.941 - 0.970] | 0.968 [0.954 - 0.979] | 0.929 [0.910 - 0.947] |
| | AUROC of run #4 | 0.955 [0.937 - 0.970] | 0.953 [0.937 - 0.967] | 0.945 [0.930 - 0.960] | 0.948 [0.930 - 0.963] | 0.929 [0.910 - 0.946] |
| | AUROC of run #5 | 0.957 [0.943 - 0.971] | 0.971 [0.959 - 0.981] | 0.952 [0.937 - 0.966] | 0.947 [0.930 - 0.962] | 0.947 [0.930 - 0.963] |
| | **Mean AUROC +/- SD** | **0.960 [± 0.009]** | **0.958 [± 0.010]** | **0.953 [± 0.005]** | **0.942 [± 0.022]** | **0.937 [± 0.007]** |
| | **Median AUROC +/- IQR** | **0.957 [± 0.012]** | **0.962 [± 0.012]** | **0.956 [± 0.004]** | **0.947 [± 0.001]** | **0.940 [± 0.012]** |
| **Gastric cancer subtyping** *raw data for Figure 1E* | AUROC of run #1 | 0.785 [0.726 - 0.844] | 0.772 [0.708 - 0.830] | 0.721 [0.646 - 0.798] | 0.725[0.686 - 0.816] | 0.743 [0.668 - 0.809] |
| | AUROC of run #2 | 0.796 [0.733 - 0.858] | 0.787 [0.723 - 0.843] | 0.735 [0.660 - 0.809] | 0.558 [0.478 - 0.632] | 0.725 [0.647 - 0.792] |
| | AUROC of run #3 | 0.758 [0.690 - 0.823] | 0.745 [0.684 - 0.812] | 0.739 [0.670 - 0.809] | 0.775 [0.712 - 0.831] | 0.753 [0.685 - 0.817] |
| | AUROC of run #4 | 0.795 [0.731 - 0.858] | 0.759 [0.696 - 0.821] | 0.752 [0.679 - 0.821] | 0.688 [0.610 - 0.758] | 0.728 [0.657 - 0.797] |
| | AUROC of run #5 | 0.774 [0.708 - 0.834] | 0.779 [0.719 - 0.839] | 0.732 [0.661 - 0.800] | 0.789 [0.735 - 0.846] | 0.737 [0.662 - 0.806] |
| | **Mean AUROC +/- SD** | **0.782 [± 0.014]** | **0.768 [± 0.015]** | **0.736 [± 0.01]** | **0.712 [± 0.085]** | **0.737 [± 0.01]** |
| | **Median AUROC +/- IQR** | **0.785 [± 0.021]** | **0.772 [± 0.02]** | **0.735 [± 0.007]** | **0.752 [0.087]** | **0.737 [± 0.015]** |

**Suppl. Table 2: Baseline performance for ResNet and ViT on both classification tasks, no attack at inference.** SD = standard deviation, IQR = interquartile range. Adversarially robust training was performed using PGD attack with ε = 0.5e-2.

| | | Is the noise detectable for a human observer? | |
|---|---|---|---|
| | | undetectable | detectable |
| **PGD attack on ResNet**<br><br>*Raw data for Suppl. Figure 2A* | **Sum** | **54** | **96** |
| | **0.0 < ε < 0.1** | 30 | 0 |
| | **0.1 < ε < 0.2** | 23 | 7 |
| | **0.2 < ε < 0.3** | 1 | 29 |
| | **0.3 < ε < 0.4** | 0 | 30 |
| | **0.4 < ε < 0.5** | 0 | 30 |
| **PGD attack on ViT**<br><br>*Raw data for Suppl. Figure 2B* | **Sum** | **38** | **112** |
| | **0.0 < ε < 0.1** | 30 | 0 |
| | **0.1 < ε < 0.2** | 8 | 22 |
| | **0.2 < ε < 0.3** | 0 | 30 |
| | **0.3 < ε < 0.4** | 0 | 30 |
| | **0.4 < ε < 0.5** | 0 | 30 |

**Suppl. Table 3: Results of the blinded observer study.** The number of images classified in the blinded observer as detectable and undetectable noise for both ResNet and ViT models.

| Experimental setup | Experimental run | Attack strength at inference | | | |
|---|---|---|---|---|---|
| | | None, ε = 0.0e-3 | Low, ε = 0.25e-3 | Medium, ε = 0.75e-3 | High, ε = 1.50e-3 |
| Model: ResNet<br>Task: RCC subtyping<br>Train: normal<br>Inference: PGD attack | AUROC of run #1 | 0.967 [0.953 - 0.978] | 0.883 [0.853 - 0.908] | 0.494 [0.450 - 0.539] | 0.058 [0.043 - 0.075] |
| | AUROC of run #2 | 0.975 [0.964 - 0.984] | 0.940 [0.921 - 0.956] | 0.769 [0.729 -0.804] | 0.331 [0.291 - 0.375] |
| | AUROC of run #3 | 0.948 [0.931 - 0.965] | 0.930 [0.908 - 0.949] | 0.869 [0.838 - 0.897] | 0.734 [0.693 - 0.772] |
| | AUROC of run #4 | 0.955 [0.937 - 0.970] | 0.921 [0.896 - 0.942] | 0.804 [0.767 - 0.837] | 0.499 [0.455 - 0.543] |
| | AUROC of run #5 | 0.957 [0.943 - 0.971] | 0.923 [0.901 - 0.943] | 0.809 [0.772 - 0.842] | 0.525 [0.476 - 0.570] |
| | **Mean AUROC +/- SD** | **0.960 [± 0.009]** | **0.919 [±0.019]** | **0.749 [± 0.131]** | **0.429 [± 0.226]** |
| | **Median AUROC +/- IQR** | **0.957 [± 0.012]** | **0.923 [± 0.009]** | **0.804 [± 0.04]** | **0.499 [± 0.194]** |
| Model: ViT<br>Task: RCC subtyping<br>Train: normal<br>Inference: PGD attack | AUROC of run #1 | 0.965 [0.952 - 0.976] | 0.954 [0.939 - 0.968] | 0.925 [ 0.903 - 0.944] | 0.861 [0.829 - 0.889] |
| | AUROC of run #2 | 0.962 [0.949 - 0.973] | 0.947 [0.930 - 0.962] | 0.911 [0.888 - 0.932] | 0.831 [0.798 - 0.860] |
| | AUROC of run #3 | 0.941 [0.923- 0.958] | 0.929 [0.911 - 0.948] | 0.900 [0.876 - 0.923] | 0.844 [0.813 - 0.874] |
| | AUROC of run #4 | 0.953 [0.937 - 0.967] | 0.933 [0.914 - 0.950] | 0.830 [0.855 - 0.906] | 0.768 [0.729 - 0.801] |
| | AUROC of run #5 | 0.971 [0.959 - 0.981] | 0.958 [0.943 - 0.970] | 0.922 [ 0.901 - 0.940] | 0.833 [0.801 - 0.863] |
| | **Mean AUROC +/- SD** | **0.958 [± 0.01]** | **0.944 [± 0.011]** | **0.908 [± 0.015]** | **0.827 [± 0.032]** |
| | **Median AUROC +/- IQR** | **0.962 [± 0.012]** | **0.947 [± 0.021]** | **0.911 [± 0.022]** | **0.833 [± 0.013]** |
| | **p versus ResNet** | **p = 0.98, t = 0.28** | **p = 0.06, t =-2.21** | **p = 0.04, t = -2.41** | **p = 0.01, t = -3.49** |

**Suppl. Table 4: Performance of ResNet and ViT on the RCC subtyping task, attacked with PGD at inference.**

| Experimental setup | Experimental run | Attack strength at inference | | | |
|---|---|---|---|---|---|
| | | **None, ε = 0.0e-3** | **Low, ε = 0.25e-3** | **Medium, ε = 0.75e-3** | **High, ε = 1.50e-3** |
| Model: ResNet | AUROC of run #1 | 0.967 [0.953 - 0.978] | 0.932 [0.909 - 0.950] | 0.825 [0.791 - 0.856] | 0.564 [ 0.519 - 0.609] |
| Task: RCC subtyping | AUROC of run #2 | 0.975 [0.964 - 0.984] | 0.960 [ 0.945 - 0.972] | 0.917 [ 0.894 - 0.937] | 0.815 [ 0.781 - 0.846] |
| | AUROC of run #3 | 0.948 [0.931 - 0.965] | 0.940 [0.921 - 0.958] | 0.917 [0.894 - 0.939] | 0.876 [0.847 - 0.903] |
| Train: normal | AUROC of run #4 | 0.955 [0.937 - 0.970] | 0.940 [0.918 - 0.958] | 0.901 [0.873 - 0.925] | 0.820 [0.784 - 0.852] |
| Inference: FGSM attack | AUROC of run #5 | 0.957 [0.943 - 0.971] | 0.943 [ 0.925 - 0.959] | 0.906 [ 0.881 - 0.929] | 0.835 [0.800 - 0.866] |
| | **Mean AUROC +/- SD** | **0.960 [± 0.009]** | **0.943 [± 0.009]** | **0.893 [± 0.035]** | **0.782 [± 0.111]** |
| | **Median AUROC +/- IQR** | **0.957 [± 0.012]** | **0.940 [± 0.003]** | **0.906 [± 0.016]** | **0.820 [± 0.020]** |
| Model: ViT | AUROC of run #1 | 0.965 [0.952 - 0.976] | 0.960 [0.946 - 0.972] | 0.949 [0.932 - 0.964] | 0.929 [0.908 - 0.947] |
| Task: RCC subtyping | AUROC of run #2 | 0.962 [0.949 - 0.973] | 0.956 [0.942-0.968] | 0.940 [0.924 - 0.955] | 0.914 [0.892 - 0.933] |
| | AUROC of run #3 | 0.941 [0.923- 0.958] | 0.936 [0.915 - 0.952] | 0.924 [ß-901 - 0.942] | 0.902 [0.877 - 0.924] |
| Train: normal | AUROC of run #4 | 0.953 [0.937 - 0.967] | 0.944 [0.926 - 0.960] | 0.923 [0.902 - 0.943] | 0.887 [0.862 - 0.912] |
| Inference: FGSM attack | AUROC of run #5 | 0.971 [0.959 - 0.981] | 0.964 [0.950 - 0.976] | 0.951 [0.934 - 0.965] | 0.926 [0.906 - 0.944] |
| | **Mean AUROC +/- SD** | **0.958 [± 0.010]** | **0.952 [± 0.010]** | **0.937 [± 0.012]** | **0.912 [± 0.016]** |
| | **Median AUROC +/- IQR** | **0.962 [± 0.012]** | **0.956 [± 0.016]** | **0.940 [± 0.025]** | **0.914 [± 0.024]** |
| | **p versus ResNet** | **p = 0.78, t = 0.28** | **p = 0.23, t = -1.29** | **p = 0.04, t = -2.41** | **p = 0.05, t = -2.31** |

**Suppl. Table 5: Performance of ResNet and ViT on the RCC subtyping task, attacked with FGSM at inference.**

| Experimental setup | Experimental run | Attack strength at inference | | | |
|---|---|---|---|---|---|
| | | None, ε = 0.0e-3 | Low, ε = 0.25e-3 | Medium, ε = 0.75e-3 | High, ε = 1.50e-3 |
| Model: ResNet<br><br>Task: Gastric cancer subtyping<br><br>Train: normal<br><br>Inference: PGD attack | AUROC of run #1 | 0.785 [0.726 - 0.844] | 0.326 [0.259 - 0.398] | 0.006 [0.002 - 0.014] | 0.000 [0.000 - 0.000] |
| | AUROC of run #2 | 0.796 [0.733 - 0.858] | 0.363 [0.292 - 0.435] | 0.007 [0.002 - 0.014] | 0.000 [0.000 - 0.000] |
| | AUROC of run #3 | 0.758 [0.690 - 0.823] | 0.519 [0.435 - 0.597] | 0.112 [0.074 - 0.153] | 0.001 [0.000 - 0.002] |
| | AUROC of run #4 | 0.795 [0.731 - 0.858] | 0.401 [0.329 - 0.477] | 0.017 [0.006 - 0.033] | 0.000 [0.000 - 0.000] |
| | AUROC of run #5 | 0.774 [0.708 - 0.834] | 0.293 [0.225 - 0.364] | 0.003 [0.000 - 0.008] | 0.000 [0.000 - 0.000] |
| | **Mean AUROC +/- SD** | **0.782 [± 0.014]** | **0.380 [± 0.078]** | **0.029 [± 0.042]** | **0.000 [± 0.000]** |
| | **Median AUROC +/- IQR** | **0.785 [± 0.021]** | **0.363 [± 0.075]** | **0.007 [± 0.011]** | **0.000 [± 0.000]** |
| Model: ViT<br><br>Task: Gastric cancer subtyping<br><br>Train: normal<br><br>Inference: PGD attack | AUROC of run #1 | 0.772 [0.708 - 0.830] | 0.676 [0.605 - 0.747] | 0.437 [0.361 - 0.513] | 0.175 [0.123 - 0.237] |
| | AUROC of run #2 | 0.787 [0.723 - 0.843] | 0.726 [0.657 - 0.791] | 0.589 [0.510 - 0.664] | 0.382 [0.305 - 0.454] |
| | AUROC of run #3 | 0.745 [0.684 - 0.812] | 0.698 [0.633 - 0.767] | 0.600 [0.527 - 0.673] | 0.446 [0.375 - 0.524] |
| | AUROC of run #4 | 0.759 [0.696 - 0.821] | 0.581 [0.506 - 0.654] | 0.255 [0.190 - 0.322] | 0.028 [0.011 - 0.05] |
| | AUROC of run #5 | 0.779 [0.719 - 0.839] | 0.729 [ 0.666 - 0.794] | 0.631 [0.560 - 0.704] | 0.459 [0.386 - 0.545] |
| | **Mean AUROC +/- SD** | **0.768 [± 0.015]** | **0.682 [± 0.054]** | **0.502 [± 0.141]** | **0.298 [± 0.169]** |
| | **Median AUROC +/- IQR** | **0.772 [± 0.02]** | **0.698 [± 0.050]** | **0.589 [± 0.163]** | **0.382 [± 0.271]** |
| | **P-value ResNet vs. ViT** | **p = 0.24, t = 1.28** | **p = 0.00, t = - 6.35** | **p = 0.00, t = - 6.45** | **p = 0.01, t = - 3.52** |

**Suppl. Table 6: Performance of ResNet and ViT on the gastric cancer subtyping task, attacked with PGD at inference.**

| Experimental setup | Experimental run | Attack strength at inference | | | |
|---|---|---|---|---|---|
| | | None, ε = 0.0e-3 | Low, ε = 0.25e-3 | Medium, ε = 0.75e-3 | High, ε = 1.50e-3 |
| Model: ResNet<br><br>Task: RCC subtyping<br><br>Train: adversarially robust training<br><br>Inference: PGD attack | AUROC of run #1 | 0.956 [0.942 - 0.969] | 0.951 [ 0.936 - 0.966] | 0.944 [0.927 - 0.959] | 0.930 [0.912 - 0.948] |
| | AUROC of run #2 | 0.958 [0.944 - 0.971] | 0.955 [0.942 - 0.969] | 0.949 [0.933 - 0.964] | 0.938 [0.921 - 0.956] |
| | AUROC of run #3 | 0.956 [0.941 - 0.970] | 0.953 [0.938 - 0.968] | 0.946 [0.930 - 0.961] | 0.933 [0.915 - 0.951] |
| | AUROC of run #4 | 0.945 [0.930 - 0.960] | 0.943 [0.927 - 0.958] | 0.937 [0.920 - 0.953] | 0.927 [0.909 - 0.945] |
| | AUROC of run #5 | 0.952 [0.937 - 0.966] | 0.947 [0.932 - 0.962] | 0.939 [0.923 - 0.956] | 0.926 [0.908 - 0.944] |
| | **Mean AUROC +/- SD** | **0.953 [± 0.005]** | **0.950 [± 0.004]** | **0.943 [± 0.004]** | **0.931 [± 0.004]** |
| | **Median AUROC +/- IQR** | **0.956 [± 0.004]** | **0.951 [± 0.006]** | **0.944 [± 0.007]** | **0.930 [± 0.006]** |
| Model: ViT<br><br>Task: RCC subtyping<br><br>Train: adversarially robust training<br><br>Inference: PGD attack | AUROC of run #1 | 0.940 [0.923 - 0.957] | 0.936 [0.17 - 0.953] | 0.927 [0.907 - 0.946] | 0.914 [0.893 - 0.935] |
| | AUROC of run #2 | 0.941 [0.924 - 0.958] | 0.936 [0.917 - 0.954] | 0.927 [0.907 - 0.946 | 0.913 [0.891 - 0.933 |
| | AUROC of run #3 | 0.929 [0.910 - 0.947] | 0.926 [0.906 - 0.943] | 0.916 [0.895 - 0.935] | 0.900 [0.878 - 0.921] |
| | AUROC of run #4 | 0.929 [0.910 - 0.946] | 0.920 [0.900 - 0.940] | 0.903 [0.881 - 0.924] | 0.872 [0.848 - 0.897] |
| | AUROC of run #5 | 0.947 [0.928 . 0.962] | 0.942 [0.925 - 0.959] | 0.932 [0.912 - 0.950] | 0.915 [0.893 - 0.936] |
| | **Mean AUROC +/- SD** | **0.937 [± 0.007]** | **0.932 [± .008]** | **0.921 [± 0.01]** | **0.903 [± 0.016]** |
| | **Median AUROC +/- IQR** | **0.940 [± 0.012]** | **0.936 [± 0.01]** | **0.927 [± 0.011]** | **0.913 [± 0.014]** |
| | **p versus ResNet** | **p = 0.01, t = 3.82** | **p = 0.00, t = 3.96** | **p = 0.00, t = 3.89** | **p = 0.01, t = 3.31** |
| Model: ResNet<br><br>Task: RCC subtyping<br><br>Train: adversarially robust DBN<br><br>Inference: PGD attack | AUROC of run #1 | 0.902 [0.880 - 0.923] | 0.919 [0.898 - 0.937] | 0.906 [0.884 - 0.96] | 0.882 [0.857 - 0.905] |
| | AUROC of run #2 | 0.947 [0.931 - 0.962] | 0.953 [0.939 - 0.966] | 0.949 [0.934 - 0.963] | 0.939 [0.922 - 0.955] |
| | AUROC of run #3 | 0.968 [0.954 - 0.979] | 0.955 [0.940 - 0.967] | 0.943 [0.925 - 0.957] | 0.921 [0.900 - 0.939] |
| | AUROC of run #4 | 0.948 [0.930 - 0.963] | 0.936 [0.917 - 0.953] | 0.905 [0.881 - 0.926] | 0.840 [0.810 - 0.869] |
| | AUROC of run #5 | 0.947 [0.930 - 0.962] | 0.959 [0.944 - 0.971] | 0.954 [0.937 - 0.967] | 0.943 [0.925 - 0. 959] |
| | **Mean AUROC +/- SD** | **0.942 [± 0.022]** | **0.944 [± 0.015]** | **0.931 [± 0.021]** | **0.905 [± 0.039]** |
| | **Median AUROC +/- IQR** | **0.947 [± 0.001]** | **0.953 [± 0.019]** | **0.943 [± 0.043]** | **0.921 [± 0.057]** |
| | **p versus ResNet** | **p = 0.35, t = 0.99** | **p = 0.51, t = 0.69** | **p = 0.32, t = 1.06** | **p = 0.23, t = 1.31** |

**Suppl. Table 7: Performance of adversarially robustly trained ResNet and ViT on the RCC subtyping task, attacked with PGD at inference.**

| Experimental setup | Experimental run | Attack strength at inference | | | |
|---|---|---|---|---|---|
| | | **None, ε = 0.0e-3** | **Low, ε = 0.25e-3** | **Medium, ε = 0.75e-3** | **High, ε = 1.50e-3** |
| Model: ResNet<br><br>Task: RCC subtyping<br><br>Train: adversarially robust training<br><br>Inference: FGSM attack | AUROC of run #1 | 0.956 [0.942 - 0.969] | 0.954 [0.939 - 0.967] | 0.950 [0.35 - 0.964] | 0.944 [0.927 - 0.959] |
| | AUROC of run #2 | 0.958 [0.944 - 0.971] | 0.957 [0.942 - 0.970] | 0.954 [0.938 - 0.968] | 0.949 [0.932 . 0.964] |
| | AUROC of run #3 | 0.956 [0.941 - 0.970] | 0.954 [0.940 - 0.969] | 0.952 [0.937 - 0.966] | 0.946 [0.930 - 0.961] |
| | AUROC of run #4 | 0.945 [0.930 - 0.960] | 0.944 [0.929 - 0.959] | 0.942 [0.926 - 0.957] | 0.937 [0.921 - 0.953] |
| | AUROC of run #5 | 0.952 [0.937 - 0.966] | 0.949 [0.935 - 0.964] | 0.945 [0.930 - 0.960] | 0.940 [0.923 - 0.955] |
| | **Mean AUROC +/- SD** | **0.953 [± 0.005]** | **0.952 [± 0.005]** | **0.949 [± 0.004]** | **0.943 [± 0.004]** |
| | **Median AUROC +/- IQR** | **0.956 [± 0.004]** | **0.954 [± 0.005]** | **0.950 [± 0.007]** | **0.944 [± 0.006]** |
| Model: ViT<br><br>Task: RCC subtyping<br><br>Train: adversarially robust training<br><br>Inference: FGSM attack | AUROC of run #1 | 0.940 [0.923 - 0.957] | 0.938 [0.920 - 0.955] | 0.933 [0.913 - 0.951] | 0.928 [0.907 - 0.946] |
| | AUROC of run #2 | 0.941 [0.924 - 0.958] | 0.939 [0.920 - 0.955] | 0.934 [0.915 - 0.951] | 0.928 [0.908 - 0.946] |
| | AUROC of run #3 | 0.929 [0.910 - 0.947] | 0.928 [0.908 - 0.946] | 0.923 [0.904 - 0.941] | 0.916 [0.894 - 0.935] |
| | AUROC of run #4 | 0.929 [0.910 - 0.946] | 0.925 [0.904 - 0.945] | 0.916 [0.895 - 0.938] | 0.905 [0.882 - 0.928] |
| | AUROC of run #5 | 0.947 [ 0.928 - 0.962] | 0.944 [0.926 - 0.960] | 0.940 [0.921 - 0.957] | 0.934 [0.913 - 0.951] |
| | **Mean AUROC +/- SD** | **0.937 [± 0.007]** | **0.935 [± 0007]** | **0.929 [± 0.009]** | **0.922 [± 0.010]** |
| | **Median AUROC +/- IQR** | **0.940 [± 0.012]** | **0.938 [± 0.011]** | **0.933 [± 0.011]** | **0.928 [± 0.012]** |
| | **p versus ResNet** | **p = 0.01, t = 3.82** | **p = 0.00, t = 3.96** | **p = 0.00, t = 4.02** | **p = 0.01, t = 3.74** |
| Model: ResNet<br><br>Task: RCC subtyping<br><br>Train: adversarially robust DBN<br><br>Inference: FGSM attack | AUROC of run #1 | 0.902 [0.880 - 0.923] | 0.923 [0.903 - 0.941] | 0.916 [0.895 - 0.934] | 0.906 [0.884 - 0.926] |
| | AUROC of run #2 | 0.947 [0.931 - 0.962] | 0.955 [0.941 - 0.968] | 0.952 [0.938 - 0.966] | 0.949 [0.934 - 0.963] |
| | AUROC of run #3 | 0.968 [0.954 - 0.979] | 0.957 [0.943 - 0.969] | 0.952 [0.937 - 0.965] | 0.943 [0.927 - 0.958] |
| | AUROC of run #4 | 0.948 [0.930 - 0.963] | 0.942 [0.922 - 0.959] | 0.931 [0.909 - 0.949] | 0.907 [0.882 - 0.929] |
| | AUROC of run #5 | 0.947 [0.930 - 0.962] | 0.960 [0.946 - 0.972] | 0.958 [0.944 - 0.969] | 0.954 [0.939 - 0.966] |
| | **Mean AUROC +/- SD** | **0.942 [± 0.022]** | **0.947 [± 0.014]** | **0.942 [± 0.016]** | **0.932 [± 0.021]** |
| | **Median AUROC +/- IQR** | **0.947 [± 0.001]** | **0.955 [± 0.015]** | **0.952 [± 0.021]** | **0.943 [± 0.042]** |
| | **p versus ResNet** | **p = 0.35, t = 0.99** | **p = 0.58, t = 0.58** | **p = 0.43, t = 0.83** | **p = 0.32, t = 1.07** |

**Suppl. Table 8: Performance of adversarially robustly trained ResNet and ViT on the RCC subtyping task, attacked with FGSM at inference.**

| Experimental setup | Experimental run | Attack strength at inference | | | |
|---|---|---|---|---|---|
| | | **None, ε = 0.0e-3** | **Low, ε = 0.25e-3** | **Medium, ε = 0.75e-3** | **High, ε = 1.50e-3** |
| Model: ResNet<br><br>Task: Gastric cancer subtyping<br><br>Train: adversarially robust training<br><br>Inference: PGD attack | AUROC of run #1 | 0.721 [0.646 - 0.798] | 0.710 [0.635 - 0.787] | 0.680 [0.605 - 0.757] | 0.645 [0.567 - 0.726] |
| | AUROC of run #2 | 0.735 [0.660 - 0.809] | 0.718 [0.641 - 0.795] | 0.689 [0.609 - 0.770] | 0.644 [0.566 - 0.727] |
| | AUROC of run #3 | 0.739 [0.670 - 0.809] | 0.726 [0.656 - 0.797] | 0.697 [0.626 - 0.771] | 0.662 [0.588 - 0.740] |
| | AUROC of run #4 | 0.752 [0.679 - 0.821] | 0.742 [0.669 - 0.813] | 0.721 [0.644- 0.796] | 0.687 [0.609 - 0.764] |
| | AUROC of run #5 | 0.732 [0.661 - 0.800] | 0.723 [0.651 - 0.793] | 0.704 [0.632 - 0.776] | 0.674 [0.600 - 0.748] |
| | **Mean AUROC +/- SD** | **0.736 [± 0.01]** | **0.724 [± 0.011]** | **0.698 [± 0.014]** | **0.662 [± 0.017]** |
| | **Median AUROC +/- IQR** | **0.735 [± 0.007]** | **0.723 [± 0.008]** | **0.697 [± 0.015]** | **0.662 [± 0.029]** |
| Model: ViT<br><br>Task: Gastric cancer subtyping<br><br>Train: adversarially robust training<br><br>Inference: PGD attack | AUROC of run #1 | 0.743 [0.668 - 0.809] | 0.729 [0.653 - 0.795] | 0.697 [0.623 - 0.764] | 0.648 [0.570 - 0.721] |
| | AUROC of run #2 | 0.725 [0.647 - 0.792] | 0.713 [0.635 - 0.782] | 0.686 [0.607 - 0.758] | 0.647 [0.567 - 0.723] |
| | AUROC of run #3 | 0.753 [0.685 - 0.817] | 0.738 [0.670 - 0.805] | 0.713 [0.643 - 0.782] | 0.669 [0.595 - 0.742] |
| | AUROC of run #4 | 0.728 [0.657 - 0.797] | 0.715 [0.644 - 0.786] | 0.688 [0.613 - 0.761] | 0.651 [0.575 - 0.729] |
| | AUROC of run #5 | 0.737 [0.662 - 0.806] | 0.730 [0.656 - 0.799] | 0.704 [0.628 - 0.778] | 0.669 [0.591 - 0.746] |
| | **Mean AUROC +/- SD** | **0.737 [± 0.01]** | **0.725 [± 0 .010]** | **0.698 [± 0.010]** | **0.657 [± 0.01]** |
| | **Median AUROC +/- IQR** | **0.737 [± 0.015]** | **0.729 [± 0.015]** | **0.697 [± 0.016]** | **0.651 [± 0.021]** |
| | **P-value ResNet vs. ViT** | **p = 0.85, t = - 0.2** | **p = 0.87, t = - 0.17** | **p = 0.95, t = 0.07** | **p = 0.58, t = 0.58** |

**Suppl. Table 9: Performance of adversarially robustly trained ResNet and ViT on the gastric cancer subtyping task, attacked with PGD at inference.**

| Metric | Task | Class | Original images | | | Attacked images | | |
|---|---|---|---|---|---|---|---|---|
| | | | ResNet | ViT | *Ratio ViT / ResNet* | ResNet | ViT | *Ratio ViT / ResNet* |
| **Spread within the class** (distance from points to center)<br><br>*lower is better* | RCC | ccRCC | 0.158 | **0.056** | *0.356* | 0.120 | **0.051** | *0.425* |
| | | chRCC | **0.057** | 0.064 | *1.123* | 0.123 | **0.115** | *0.935* |
| | | papRCC | 0.171 | **0.043** | *0.251* | 0.169 | **0.087** | *0.515* |
| | Gastric cancer | diffuse | 0.214 | **0.206** | *0.963* | **0.174** | 0.266 | *1.529* |
| | | intestinal | 0.216 | **0.175** | *0.810* | **0.184** | 0.260 | *1.413* |
| **Spread between classes** (distance from center to center)<br><br>*higher is better* | RCC | ccRCC to chRCC | 0.478 | **0.899** | *1.860* | 0.459 | **0.842** | *1.834* |
| | | ccRCC to papRCC | 0.560 | **0.922** | *1.646* | 0.457 | **0.896** | *1.961* |
| | | chRCC to papRCC | 0.524 | **0.882** | *1.683* | 0.434 | **0.751** | *1.730* |
| | Gastric cancer | diffuse to intestinal | 0.502 | **0.629** | *1.253* | 0.295 | **0.634** | *2.149* |

**Suppl. Table 10**. **Spread of data points within classes, and distance between classes in the latent space, related to Figure 5A-B**. The average Euclidean distance of the points within each cluster to its center for features extracted from the original images. Also it reports the distance between the center of 3 clusters. For normal images and perturbed images (PGD attack with ε of 0.05). In each pairwise comparison, the better value is printed bold.

#### Supplementary Figures

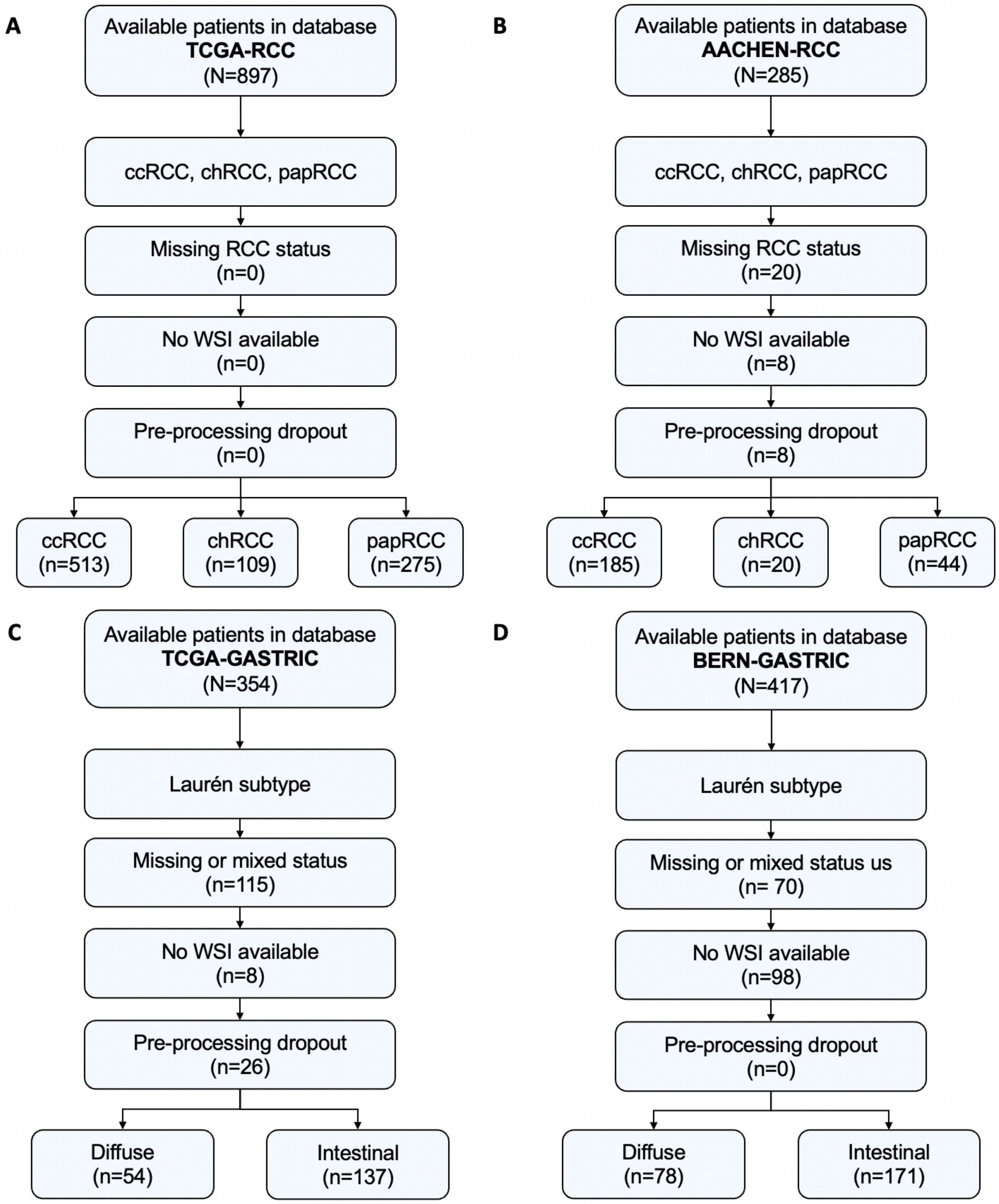


**Suppl. Figure 1:** CONSORT charts for all cohorts in this study. **(A)** TCGA-RCC, **(B)** AACHEN-RCC, **(C)** TCGA-GASTRIC, **(D)** BERN-GASTRIC. N = total patient number, n = patient number of subset.

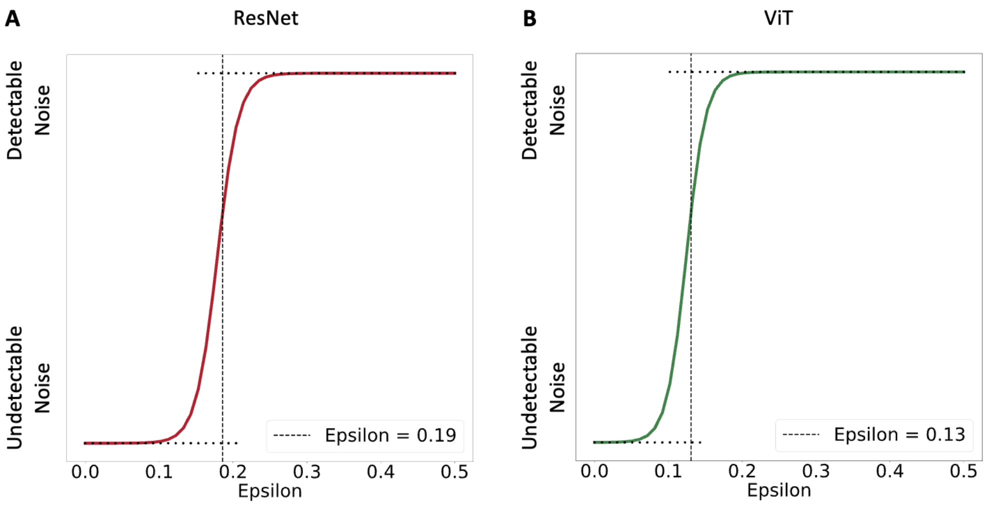


**Suppl. Figure 2: Determining the observable threshold for adversarial attacks. (A)** In a blinded user study, 150 images with different amounts of noise were assessed. The results were analyzed with a logistic regression. For ResNet, the observable threshold for adversarial noise was ε=0.19. **(B)** For ViT, the observable threshold for adversarial noise was ε=0.13. Raw data for this figure is given in Suppl. Table 3.

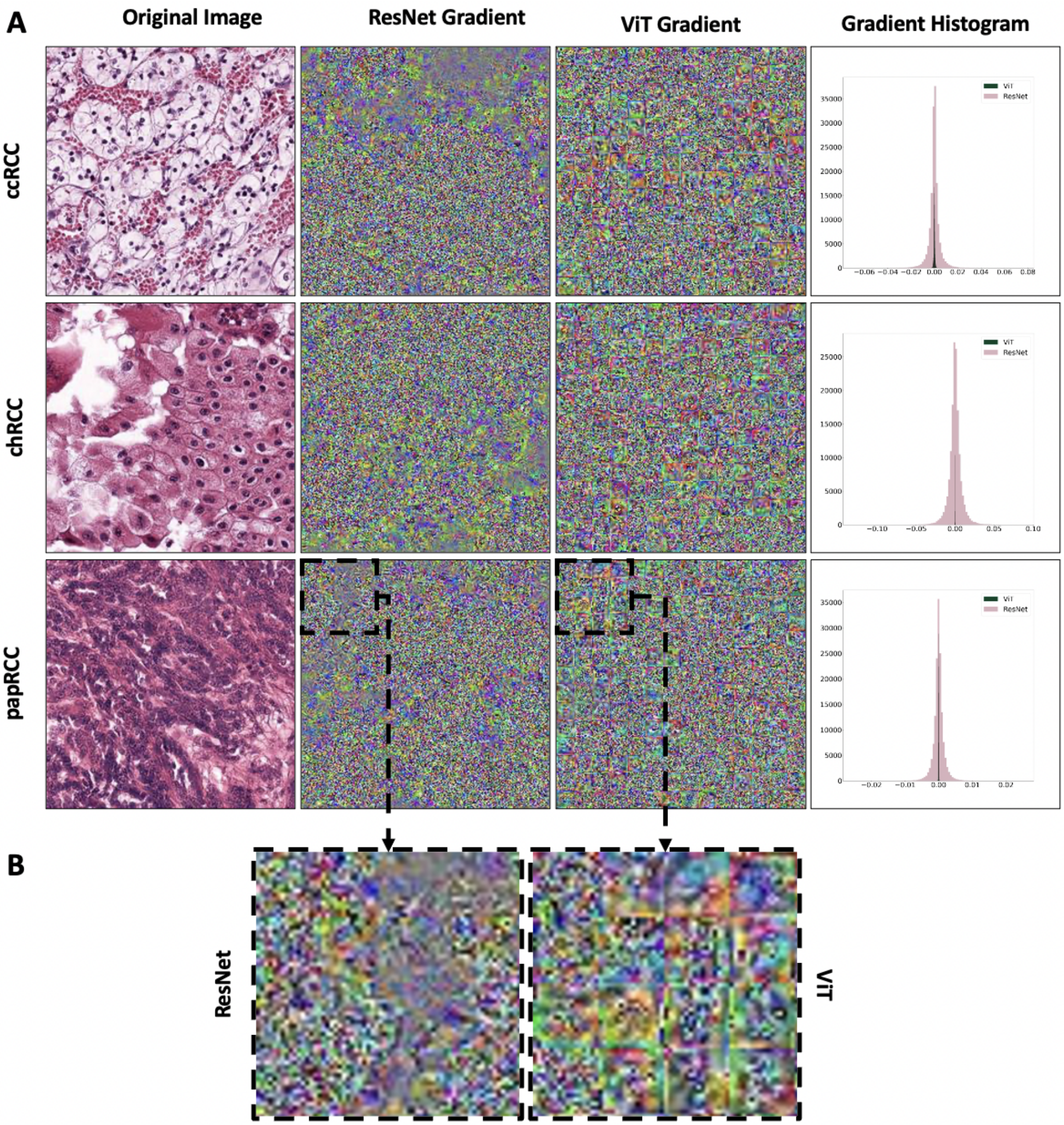


**Suppl. Figure 3: Visualization of PGD gradients for representative images. (A)** One representative image for each class in the RCC subtyping task was used to visualize the PGD gradients obtained with a white-box attack on a ResNet and a ViT model. **(B)** The enlarged detail shows that the adversarial noise follows the patch structure of the Transformer.

### Supplementary Methods

#### Deep Learning models

We used two types of Deep Learning models: Residual neural networks (ResNets) and Vision Transformers (ViTs).

ResNets were developed in 2015 to overcome the vanishing gradient effect problem when training deep neural networks. In addition to possessing all common elements of typical convolutional neural networks such as convolution, pooling, activation and fully connected layers, this architecture also uses identity connection between residual blocks to add the output from the previous layer to the layer ahead of it. Resnet architectures are currently the state of the art in computational pathology.[23]. We used ResNet50 pre-trained on ImageNet as the initial model. The first $50\%$ of the layers were frozen and only the last $50\%$ of them were fine-tuned during the training process. The input for the ResNet50 are tiles with a size of $(224 \times 224 \times 3)$ . The output for each image after a softmax layer is the prediction scores belonging to each target class.

ViTs were introduced in 2021 for image classification tasks after transformers demonstrated enormous success in natural language processing tasks. ViTs work by splitting the input image into small patches and then by using linear projection, creating patch-embeddings from the flattened patches. In addition to this, it adds positional embedding into the patch-embeddings and feeds this sequence as an input into the standard transformer encoder. As a vision transformer model, we used the 'B_16_imagenet1k' model pre-trained on ImageNet with a minor modification of the input size $(224 \times 224 \times 3)$ from the pytorch_pretrained_vit Python package. As with ResNet, the output of ViT for each tile is the prediction scores for each class.

#### Hyperparameters

To find the optimum learning rate for the training we used the torch_lr_finder Python package with an initial learning rate of $10^{-5}$ and weight_decay of $10^{-2}$ for 100 iterations with an end learning rate of 100. We used Adam optimizer during training with the calculated $\frac{optimum\ learning\ rate}{10}$ and a weight-decay of $10^{-2}$. The defined loss function for all experiments was cross-entropy. For all the models (ResNet, ViT, and DBN), we resize the input images to $(224 \times 224 \times 3)$ in order to have a fair comparison between different architectures. In each training run, we randomly selected $20\%$ of the patients for the validation set to stop the training using early stopping with the minimum training epoch of 10 and the patience of 5.

#### Types of adversarial attacks

Adversarial attacks can be both white-box attacks in which the attacker has access to the model's parameters, or black box attacks in which the attacker has no access to the parameters and it uses either a different model or no model to perturb the images. Additionally, the attacks can be targeted attacks to force the model to output a specific class prediction for an input image or non-target attacks in which general misclassification is the primary purpose. In this study, we use three untargeted white-box attacks:

1. **Fast Gradient Sign Method (FGSM)**.[36–38] This is a white-box attack which generates an adversarial image by changing the pixel magnitudes in the direction of the gradient. This attack is a single-step attack and is therefore very efficient in terms of computation time. Default parameters are iteration= 1, objective function = infinity norm.
2. **Projected Gradient Descent (PGD)**. This is a multi-step attack based on FGSM and is considered as one of the most powerful and complete white-box adversaries [39]. PGD generates perturbations to maximize the value of loss function while constraining the changes in a specified value, defined as epsilon ($\epsilon$). Because of its iterative nature, PGD is more time consuming. Models that are robust to PGD are always robust against other forms of gradient-based attacks [54,55]. Default parameters are iteration= 10, objective function = infinity norm. For the example, in **Figure 1C**, these attack parameters were used: $\varepsilon = 0.005$, alpha = 0.0025. For all other attacks, three levels of $\varepsilon$ were pre-defined: low ($0.25 \times 10^{-3}$), medium ($0.75 \times 10^{-3}$), and high ($1.5 \times 10^{-3}$).
3. **Fast Adaptive boundary (FAB)**.[40] This white-box attack searches for the minimum perturbation which is required to change the class of an input image. Due to the architecture of FAB, to check the success rate of models, it doesn't require repeating the attack for different epsilon values. This stands in contrast to PGD, as described above. As such, it provides a more complete picture of the robustness of a specific model while additionally requiring more computational time in comparison to other white-box attacks.

In addition, we used an untargeted black-box attack:

4. **Square attack.**[41] Unlike the white-box attacks described above, a square attack is a query-efficient black-box attack which does not rely on local gradient information of the specific model.[41] This iterative black-box adversary places square-shaped updates at random positions on the input image and searches for increases in the loss function in each iteration. While square attacks have the novelty of being significantly different in approach to most other attacks, they usually have a lower success rate and are more computationally expensive than white-box attacks.[52,56,57]

#### Attack mitigation strategies

We used two attack mitigation strategies: adversarially robust training and dual batch normalization.

To robustly train ResNet and ViT, we attacked the images with a PGD white-box attack (ε = 0.005, alpha = 0.0025 and the objective function = infinity norm) just before feeding them to the models during the training and subsequently calculated the loss function based on the prediction of models for perturbed images. We evaluated adversarially pre-trained models in concordant attacks (e.g. train with PGD, test with PGD) and discordant attacks (e.g. train with PGD, test with FGSM).

The dual batch adversarial robust model was introduced to the medical field by Han et al. in 2021[35] . This model is a modified ResNet50 with two batch norm layers, one for the standard input and the other for the adversarially perturbed inputs. We use the same hyperparameters for adversarial attacks in this training. The total loss is the sum of the normal loss function for the standard inputs and the loss function for the perturbed images for the same label.

#### Hardware

All experiments were run on local computer workstations with Nvidia RTX A6000 and Quadro RTX 8000 graphics processing units (GPUs).
